# Transient effects in corticospinal and reticulospinal tract excitability induced by motor skill and isometric resistance training

**DOI:** 10.1101/2025.05.21.655351

**Authors:** Rachel Hawthorn, Natalie Phelps, Carolyn Atkinson, Rodolfo Keesey, Zachary Seitz, Haolin Nie, Ismael Seáñez

**Author notes:** **Corresponding author: Ismael Seáñez, PhD,**.

## Abstract

Motor skill and resistance training are commonly used in rehabilitation to enhance neural plasticity. Nonetheless, how each modality impacts the excitability of corticospinal and reticulospinal pathways controlling the lower limb remains poorly understood. Here, we tested how single 30-minute sessions of cue-paced motor skill and isometric resistance training modulate corticospinal, reticulospinal, and spinal excitability in unimpaired adults (N = 23). Using motor-evoked potentials via transcranial magnetic stimulation, we found that both training types increased corticospinal excitability, with more substantial effects following motor skill training. In contrast, reticulospinal tract excitability—assessed by StartReact responses—and spinal excitability—assessed by H/M ratios, F-wave response amplitude, and persistence—remained largely unchanged. These results suggest that short-term training selectively enhances corticospinal tract excitability without a measurable impact on spinal or reticulospinal circuits. This pathway-specific response may inform strategies for targeting neural plasticity in rehabilitation.

## INTRODUCTION

Spinal cord injuries (SCI) cause long-lasting damage to descending motor pathways, resulting in impaired motor function and increased spasticity^1,2^. The activity-dependent reorganization of neural pathways observed by emerging technologies for neurorehabilitation^3–5^ suggests a paradigm shift from compensation to recovery-based interventions for SCI^6^. Therefore, identifying the neural pathways that are prone to plasticity via activity-based interventions can promote the development of new rehabilitation strategies that specifically target these circuits to enhance and accelerate recovery after SCI.

The corticospinal and reticulospinal tracts are two major tracts with descending projections onto spinal motoneurons and interneurons in mammals^7–9^. The corticospinal tract has been shown to contribute to the voluntary control of walking^10,11^ and distal movements^12^. While corticospinal tract excitability has been shown to increase after motor skill training^13^, it appears to remain unchanged after resistance^14^, balance^15^, and exercise-induced fatigue training^16^. However, a recent study has shown that synchronizing strength training with an external cue, such as a metronome, may add a skill or complexity component to the training, thereby increasing corticospinal tract excitability^17^. The reticulospinal tract has been shown to contribute to postural adjustments^20^, gross hand function^21^, selecting the appropriate force level during movements^22,23^, modulating flexor and extensor motor neurons to maintain balance^24,25^, and performing bimanual tasks^26^. The spinal cord serves as the integrative center where corticospinal and reticulospinal projections interact with spinal circuits controlling movement^27–30^. However, studies directly comparing training-induced changes in the corticospinal, spinal, and reticulospinal tracts after similar activity-based training are lacking, limiting our understanding of where neural adaptations occur and which motor circuits are most responsive to different training modalities.

In this study, we investigated the effect of single sessions of cue-paced motor skill and cue-paced isometric resistance training on changes in corticospinal, spinal, and reticulospinal tract excitability for lower limb muscles in twenty-three unimpaired individuals. To quantify short-term changes in corticospinal, spinal motoneuron, and reticulospinal tract excitability, we assessed multiple neurophysiological responses. Corticospinal tract excitability was measured using motor-evoked potentials (MEPs) elicited by transcranial magnetic stimulation (TMS)^13,31^. Spinal excitability was evaluated through H-reflexes, M-waves^32,33^ and F-waves^27,34,35^ elicited by peripheral nerve stimulation. Reticulospinal tract excitability was assessed using the StartReact response, defined as a shortening in reaction time following a startling auditory stimulus that engages the reticulospinal tract^36^, with all measures collected before and after a 30-minute training session. We hypothesized that single session precision-control (motor skill) training would selectively enhance corticospinal tract excitability without affecting reticulospinal excitability, whereas strength (resistance) training would enhance reticulospinal tract excitability without influencing corticospinal excitability.

We found that both cue-paced motor skill and cue-paced isometric resistance training significantly increased corticospinal tract excitability. However, increases in corticospinal tract excitability were more pronounced for motor skill training than for isometric resistance training. Although we observed a significant contribution of the reticulospinal tract before and after training, no training paradigm resulted in substantial changes in reticulospinal tract excitability.

These results suggest that short-term changes in corticospinal tract excitability can be induced after single sessions of both motor skill or resistance training when an auditory cue is provided. However, the lack of training-induced changes in the StartReact response suggests that either reticulospinal tract excitability does not change after a single 30-min session of motor skill or isometric resistance training or that the reticulospinal tract is not as involved in these tasks as in people with SCI, who have shown increased contributions of the reticulospinal tract compared to unimpaired individuals^37^.

Diversity in rehabilitation paradigms will be crucial to promoting tract-specific neuroplasticity via activity-based training and emerging neuromodulation strategies^3–5,38–40^. For example, an individual with spared corticospinal tract connections may benefit from the training of dexterous tasks and distal movements^12,13,37^, whereas residual reticulospinal tract inputs may dictate the training of postural balance, strength production, and bilateral tasks^20,22,26^. Our study provides an experimental framework that can be utilized to probe how these circuits evolve after different types, durations, and intensities of training for neurorehabilitation.

## RESULTS

Thirty-two unimpaired participants provided informed consent to participate in this study, which was reviewed and approved by Washington University in St. Louis’ Institutional Review Board. The study design consisted of a randomized crossover trial with parallel group allocation. Participants were divided into two groups to assess the excitability of different pathways. Group A underwent evaluations of corticospinal and spinal excitability using transcranial magnetic stimulation (TMS) and peripheral nerve stimulation (PNS) of the posterior tibial and common peroneal nerves, respectively^27,41,42^ (**Figure 1a**). Group B underwent reticulospinal tract excitability evaluations using a StartReact evaluation^36^ (**Figure 1b**). Both groups participated in a two-arm randomized crossover design, where they performed a 30-minute motor skill training session using a novel body-machine interface (BoMI) and a 30-minute isometric resistance training session on separate days. Training session sequences were randomized for each participant to mitigate potential effects related to session order. Excitability measurements were conducted before and after each session.

**Figure 1.**
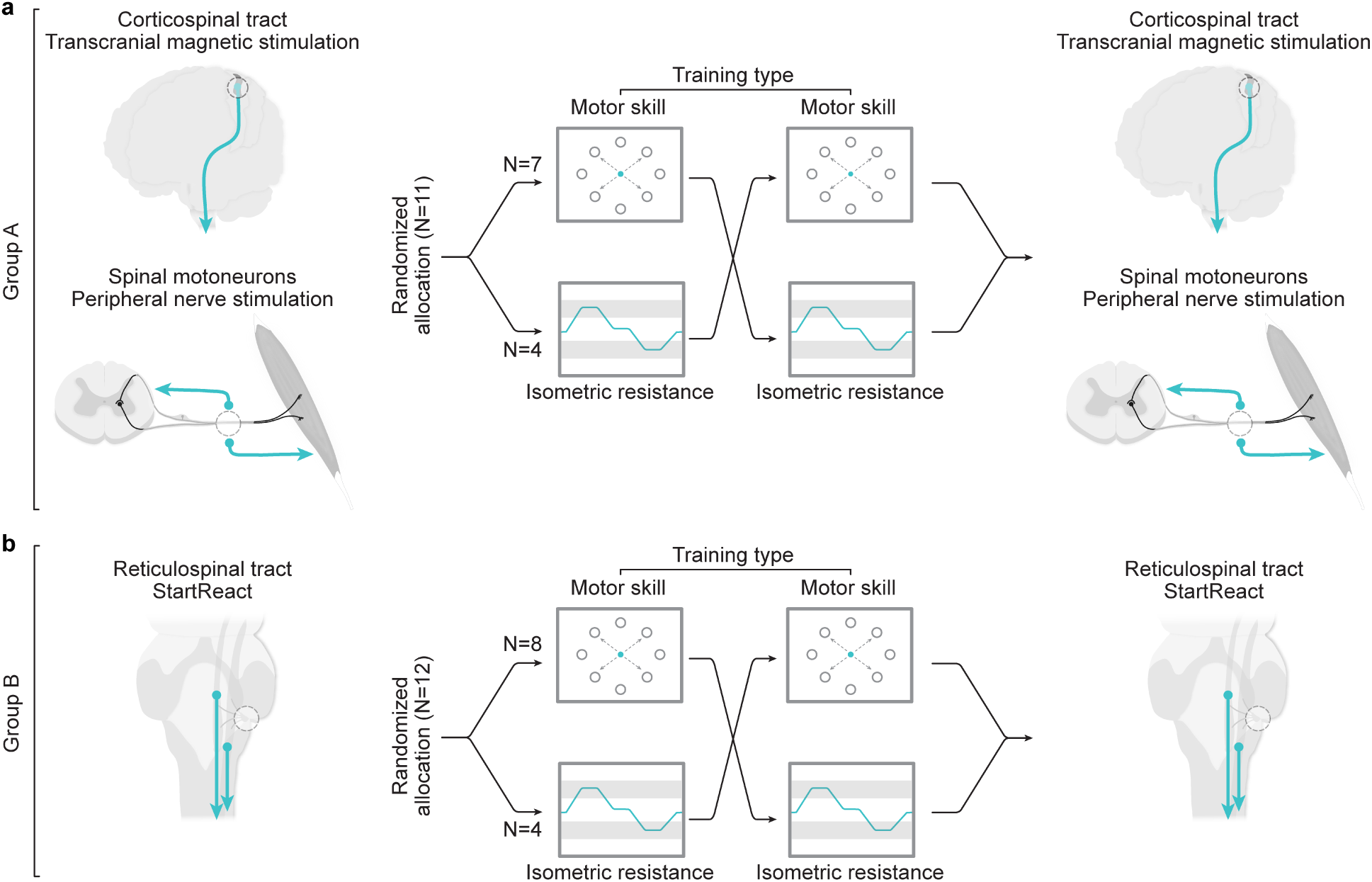
Neurophysiology experiments to evaluate cortico-reticulo-spinal excitability following motor skill and isometric resistance training. (**a**) Assessment of corticospinal and spinal excitability in Group A. Participants in Group A (N = 11) underwent transcranial magnetic stimulation (TMS) to assess corticospinal tract excitability and peripheral nerve stimulation (PNS) of the posterior tibial and common peroneal nerves to assess spinal excitability. (**b**) Assessment of reticulospinal tract excitability in Group B. Participants in Group B (N = 12) completed StartReact evaluations to measure reticulospinal tract excitability. Excitability measurements were taken immediately before and after each training session. Both groups participated in a randomized crossover trial involving a 30-minute motor skill training session using a body-machine interface (BoMI) and a 30-minute isometric resistance training session performed on separate days.

### Improvement in cursor control suggests learning of a novel motor task

A novel BoMI was used to perform leg motor skill training, requiring participants to learn precise, coordinated movements to control a cursor in a visuomotor task. Right foot and ankle movements measured with inertial measurement units (IMUs) were used to train a decoder based on principal component analysis (PCA) to convert 8-dimensional ankle movements into 2-dimensional cursor control in real-time as previously described by our group for the upper body^43^ (**Fig. 2a**). By controlling the cursor using their legs through the BoMI, participants performed five blocks of center-out reaching over the 30-minute session^44–47^. An auditory 1 Hz tone was used to incorporate a time constraint within the center-out reaching task, and participants were instructed to attempt to complete each movement at the end of the 4-second cue, moving as smoothly and efficiently as possible. Representative cursor positions and velocity profiles from the first and last blocks showed improvements in body coordination and cursor control over the training session (**Figure 2b**). Straighter cursor trajectories and smoother velocity profiles, with fewer peaks and shorter durations, characterize these improvements. Quantitative analysis of cursor control metrics revealed significant reductions in jerk (**Figure 2c**: −16.66% ± 83.98 [mean ± SD], t-statistic: 2.84, p-value: 0.010), movement time (**Figure 2d**: −18.21% ± 38.20, W: 37.00, p-value: 0.001), and path length (**Figure 2e**: −35.58% ± 39.82, W: 24.00, p-value < 0.001) following five blocks of motor skill training. These findings suggest that participants exhibited smoother, faster, and more coordinated movements during the session. Although the end-point error (distance of the cursor from the target at the 4-second mark) reduced slightly, the change was not statistically significant (−4.40% ± 28.73, W: 99.00, p-value: 0.247) (**Figure 2f**). This result is expected, as the task instructions emphasized temporal precision over spatial accuracy, thereby limiting the potential for improvements in end-point error. Together, these findings highlight the effectiveness of the motor skill training protocol in promoting motor learning of a novel task through BoMI practice^45,46,48^.

**Figure 2.**
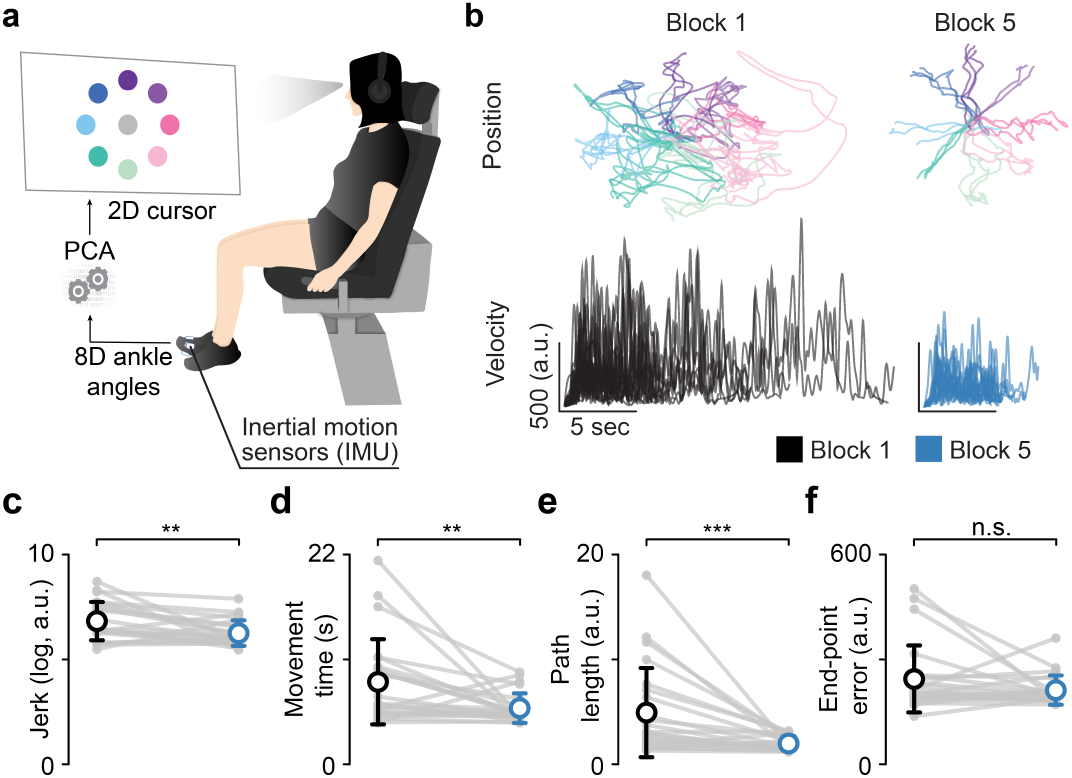
Improvements in movement coordination and control during motor skill training using an ankle-operated body-machine interface. (**a**) BoMI set up for lower-limb motor skill training. Participants used foot and ankle movements captured by IMUs to control a 2D cursor on the screen. A decoder trained with PCA translated 8-dimensional kinematic data into 2-dimensional cursor movements. (**b**) Representative cursor trajectories and velocity profiles during training. Cursor paths from an example participant (MS006) in blocks 1 and 5 illustrate straighter, more directed movements as training progressed. Average velocity traces show smoother and more coordinated movements over time, with fewer peaks and reduced variability. (**c-f**) Group-level improvements in movement quality. (**c**) Jerk. A measure of movement smoothness, defined as the rate of change of acceleration, decreased across training. (**e**) Movement time. The time taken to move from the center target to the outer target was reduced. (**e**) Path length. The total distance traveled by the cursor from the center to the target decreased across training blocks. (**f**) End-point error. The final position of the cursor at the 4-second mark, relative to the target, remained consistent. Asterisks above the group averages in (**c-f**) denote Bonferroni-corrected statistical significance from paired comparisons between performance measures of first and last training blocks: *p < 0.05, **p < 0.01, *** p < 0.001, ‘n.s.’ p > 0.05.

### Comparable neuromuscular fatigue indicates similar physical demands across protocols

The 30-minute isometric resistance protocol involved executing twelve slow-ramped isometric contractions of dorsiflexion and plantar flexion at 30% of the participants’ maximum voluntary contraction (MVC), organized into three blocks (**Figure 3a**). To ensure consistent movement timing during the ramped contractions, a 1 Hz auditory cue was employed, akin to the cue used in motor skill training. Torque traces from both the initial and final blocks exhibited stable performance throughout the training session (**Figure 3b**). To evaluate the neuromuscular fatigue and effort levels exerted by participants during both training types, maximum voluntary contractions (MVCs) were measured before and after training. Examples of maximum voluntary contractions for the tibialis anterior are illustrated in **Figure 3c**. Group analysis revealed a significant reduction in EMG amplitude following training for both motor skill (−12.97% ± 17.21, W: 31.00, p-value: 0.001, Hedges’ g: 0.285) and isometric resistance training (−6.29% ± 14.39, W: 60.00, p-value: 0.033, Hedges’ g: 0.146) (**Figure 3d**). Reductions in EMG amplitude were comparable across training modalities (W: 90.00, p-value: 0.453) (**Figure 3e**), suggesting similar levels of neuromuscular fatigue.

**Figure 3:**
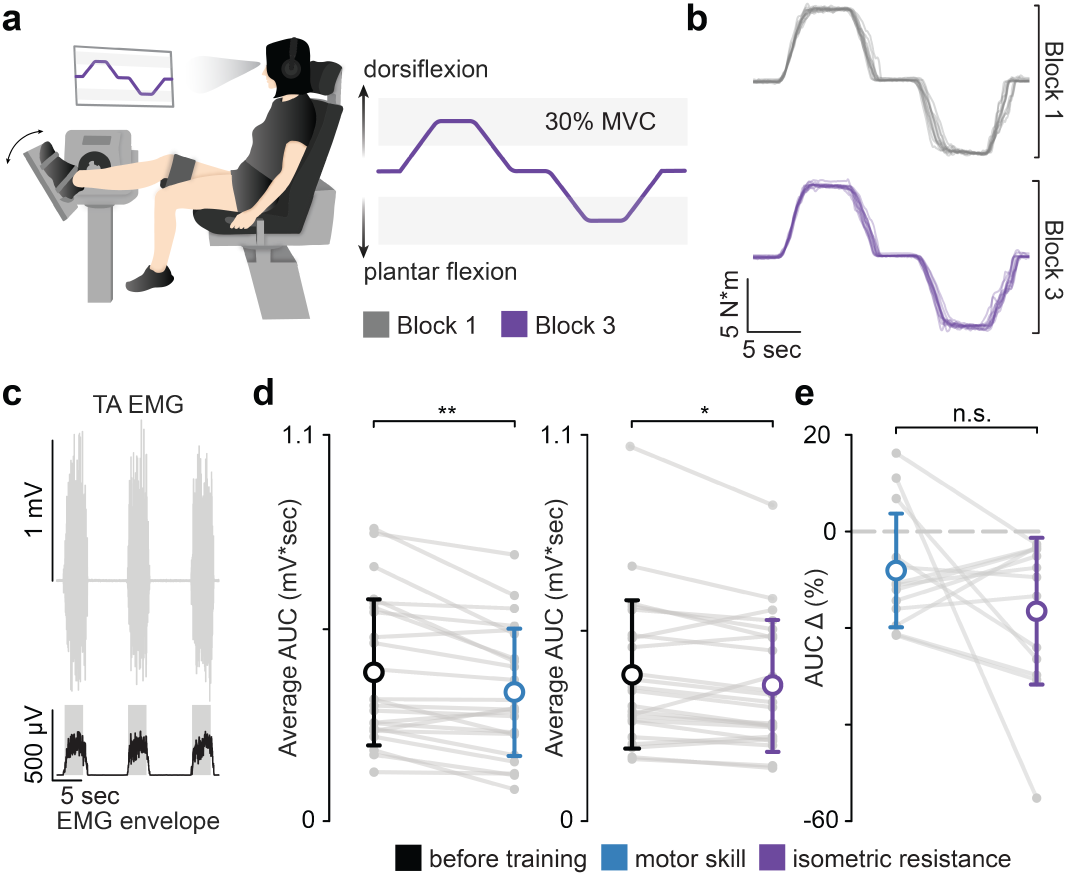
Isometric resistance and motor skill training result in similar reductions in EMG amplitude. (**a**) Isometric resistance training protocol. Participants completed three blocks of slow-ramped dorsiflexion and plantar flexion contractions at 30% of their maximum voluntary contraction (MVC), paced by a 1 Hz auditory cue to ensure consistent timing. (**b**) Representative torque traces during training. Torque recordings from participant MS005 during the first and last training blocks show consistent force generation across the session. (**c**) EMG burst and RMS traces for fatigue assessment. The top trace shows example tibialis anterior EMG bursts from participant MS018 during maximum voluntary contractions (MVCs); the bottom trace shows the root mean square (RMS) of the full-wave rectified EMG signal. The area under the RMS curve over 2.25 seconds after onset was used to quantify EMG amplitude. (**d**) Training-specific reductions in EMG amplitude. Group-level analysis revealed a decrease in EMG amplitude following both motor skill and isometric resistance training sessions. (**e**) Comparison of change in MVC amplitude across training types. Reductions in EMG amplitude were comparable between the two training protocols, indicating similar levels of neuromuscular fatigue. Asterisks above the group averages in (**d, e**) denote Bonferroni-corrected statistical significance from paired comparisons of AUC before and after individual training and between trainings: *p < 0.05, **p < 0.01, ‘n.s.’ p > 0.05.

### Motor skill training leads to higher increases in corticospinal tract excitability compared to isometric resistance training

TMS targeting the left motor cortex was used to elicit MEPs in the right tibialis anterior to assess the excitability of the corticospinal tract^13,49^ (**Figure 4a**). A custom-built NeuroNavigation software (Matlab 2020a, USA) integrated with Qualisys Track Manager (v2021.1, build 6350, Sweden) was used to identify and recall the right tibialis anterior activation hotspot at baseline and after training (**Figure 4b**). TMS at different stimulator outputs was used to compare MEP response amplitude before and after training (**Figure 4c, d**). Training-induced changes in corticospinal tract excitability were assessed by comparing percent changes in MEP amplitude at rest and during 15% maximal voluntary EMG activation of the tibialis anterior. (**Figure 4e**). Evaluation of MEPs at rest and during pre-activation provides complementary information, as rest captures baseline corticospinal tract excitability, while voluntary measures reflect pathway function during movement, albeit influenced by spinal excitability^30,50^. Results from the Generalized Linear Mixed Model (3 factors: TMS intensity [100%, 150%, 180%, 200%]; training [motor skill training vs. isometric resistance training]; evaluation condition [rest vs. pre-activation (15% dorsiflexion)]) showed a significant increase in MEP amplitudes after motor skill and resistance training (motor skill training: β = 37.21, SE = 13.20, z = 2.82, p: 0.005; isometric resistance training: β = 28.55, SE = 13.25, z = 2.15, p: 0.031, see **Supplementary Table S2** for details), suggesting that both types of training resulted in an overall increase in corticospinal tract excitability. **Figure 4f** shows the average training-induced changes in MEP response amplitudes for motor skill and resistance training during rest and pre-activation (15% dorsiflexion) evaluations. Post hoc one-sample t-tests showed that motor skill training during rest (t-statistic: 3.480, p-value: 0.005), pre-activation (W: 279.00, p-value: 0.028), and resistance training at rest (W: 299.00, p-value: 0.033) resulted in significantly greater MEP responses, while resistance training at pre-activation (t-statistic: 0.763, p-value: 1.0) caused no significant change.

**Figure 4.**
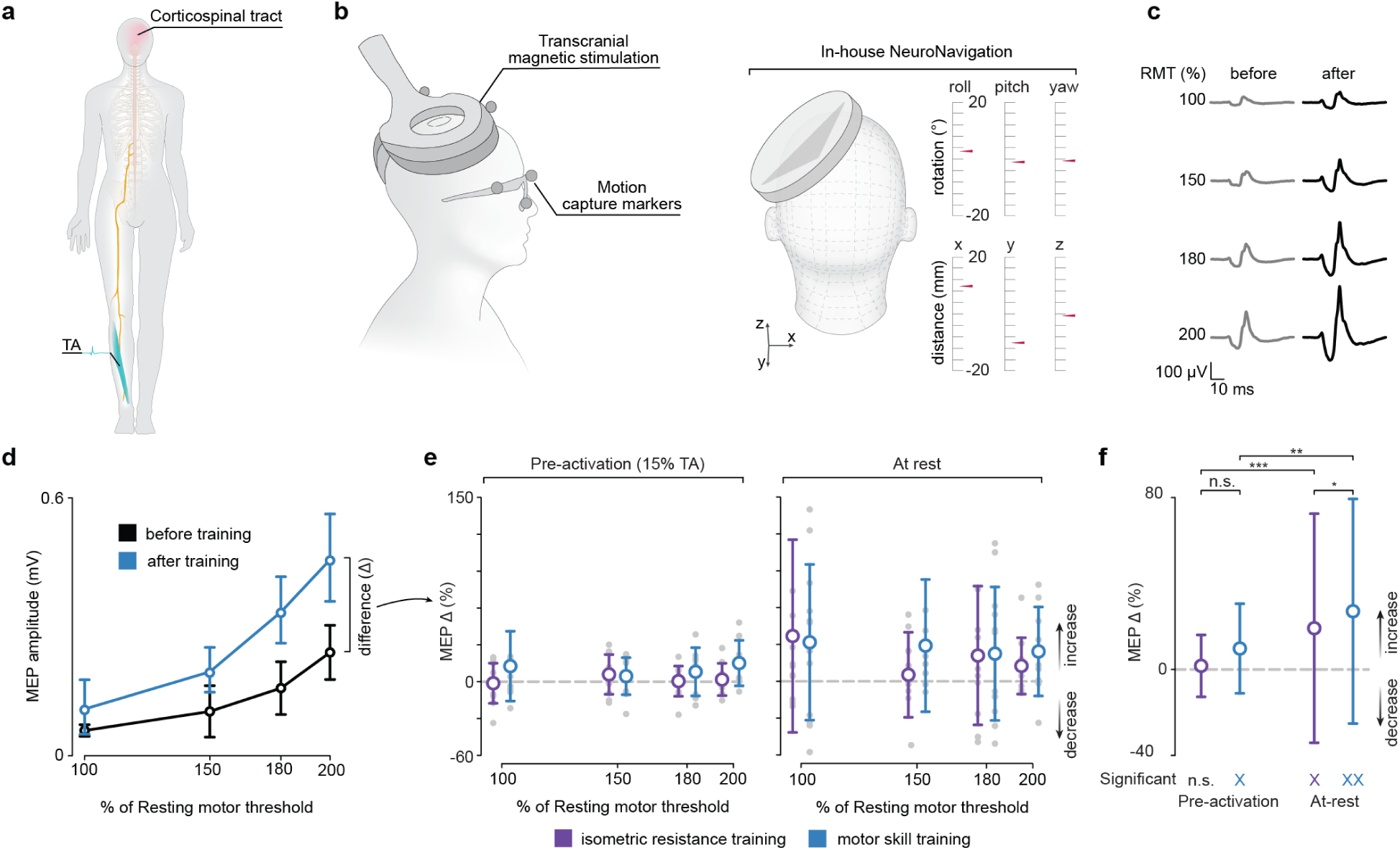
Increases in corticospinal tract excitability following motor skill and isometric resistance training measured via transcranial magnetic stimulation. (**a**) TMS was applied to the left motor cortex to elicit MEPs in the right tibialis anterior. (**b**) NeuroNavigation setup for hotspot targeting. A custom-built Neuronavigation software, using infrared camera-based motion capture markers, tracked the position and orientation of a double-cone coil to consistently target the tibialis anterior hotspot before and after each training session. (**c**) Average MEP responses at rest before and after training for a representative participant (MS004) illustrate increases in response amplitude following motor skill training. (**d**) Corresponding recruitment curve of (**c**). MEP amplitudes increased across multiple TMS intensities (100%, 150%, 180%, 200%) after training. (**e**) Percent change in MEP amplitude during pre-activation and rest conditions. The left panel shows percent changes in MEP amplitude during the pre-activation condition (15% dorsiflexion), and the right panel shows percent changes during rest. Both plots represent group-level responses across all participants following motor skill and isometric resistance training. (**f**) Training-induced changes across evaluation conditions. MEP amplitude increases following both training protocols during specific conditions. Asterisks above the bars denote post-hoc paired Bonferroni-corrected significance values for interactions between training types and evaluation conditions: *p < 0.05, **p < 0.01, *** p < 0.001, ‘n.s.’ p > 0.05. (Significant) ‘X’ markers below the plot in (**f**) indicate Bonferroni-corrected significant training effects within each condition of MEP percent change based on one-sample tests against µ = 0 (‘X’ p < 0.05, ‘XX’ p < 0.01, ‘n.s.’ p > 0.05).

We next tested how changes in MEP amplitude compared across training paradigms and evaluation conditions. Increases in MEP amplitude were significantly higher following motor skill training compared to resistance training at rest (motor skill training: 27.11% ± 52.25 increase; isometric resistance training: 19.21% ± 52.27 increase; W: 5814.00, p-value: 0.028), but not during the pre-activation condition (motor skill training: 9.75% ± 20.86 increase; isometric resistance training: 1.70% ± 14.42 increase, W: 5025.00, p-value: 0.117) (**Figure 4f**). Moreover, training-induced changes in MEP showed a higher relative change when evaluated at rest compared to the pre-activation condition (motor skill training: 16.71% ± 56.31 difference, W: 5143.00, p-value: 0.003; resistance training: 17.88% ± 51.44 difference, W: 3968.00, p-value < 0.001). These findings suggest that while both training paradigms increased corticospinal tract excitability, motor skill training resulted in a larger increase than isometric resistance training. Moreover, our results indicate that while a dorsiflexion pre-activation level of 15% can indeed increase MEP amplitudes in the tibialis anterior, changes in MEP amplitude due to training were less pronounced than when evaluated at rest.

### The majority of spinal motoneuron and monosynaptic reflex excitability measures do not show training-induced changes

The corticospinal tract contains descending projections from the motor cortex to motoneurons and interneurons in the spinal cord^8,9^. Therefore, increases in corticospinal tract excitability could be due to either an increase in the excitability of the motor cortex and its descending projections or increases in the excitability of the motoneurons projecting to the muscles and their associated synapses and interneurons. We sought to investigate whether increases in corticospinal tract excitability could be partially accounted for by increases in excitability of spinal circuits.

We used the Hoffman reflex (H-reflex), M-waves, and F-waves elicited via PNS of the mixed tibial nerve to compare the excitability of spinal neural circuits before and after training^27,41,51,52^ (**Figure 5a**). The H/M ratio is commonly used to assess monosynaptic reflex excitability (**Figure 5b**)^32,33^. One-sample tests revealed no significant changes in the H/M ratio of the TA muscle after motor skill training (25.41% ± 47.35, W: 22.00, p-value: 1.0, Hedge’s g = 0.054) or isometric resistance training (−3.72% ± 19.95, W: 20.00, p-value: 0.835, Hedge’s g = −0.138) (**Figure 5c**). Moreover, there were no significant differences in H/M ratio change between motor skill and isometric resistance training (t-statistic = 1.849, p = 0.282). These findings suggest that neither motor skill training nor isometric resistance training significantly increases the excitability of the tibialis anterior monosynaptic reflex.

**Figure 5.**
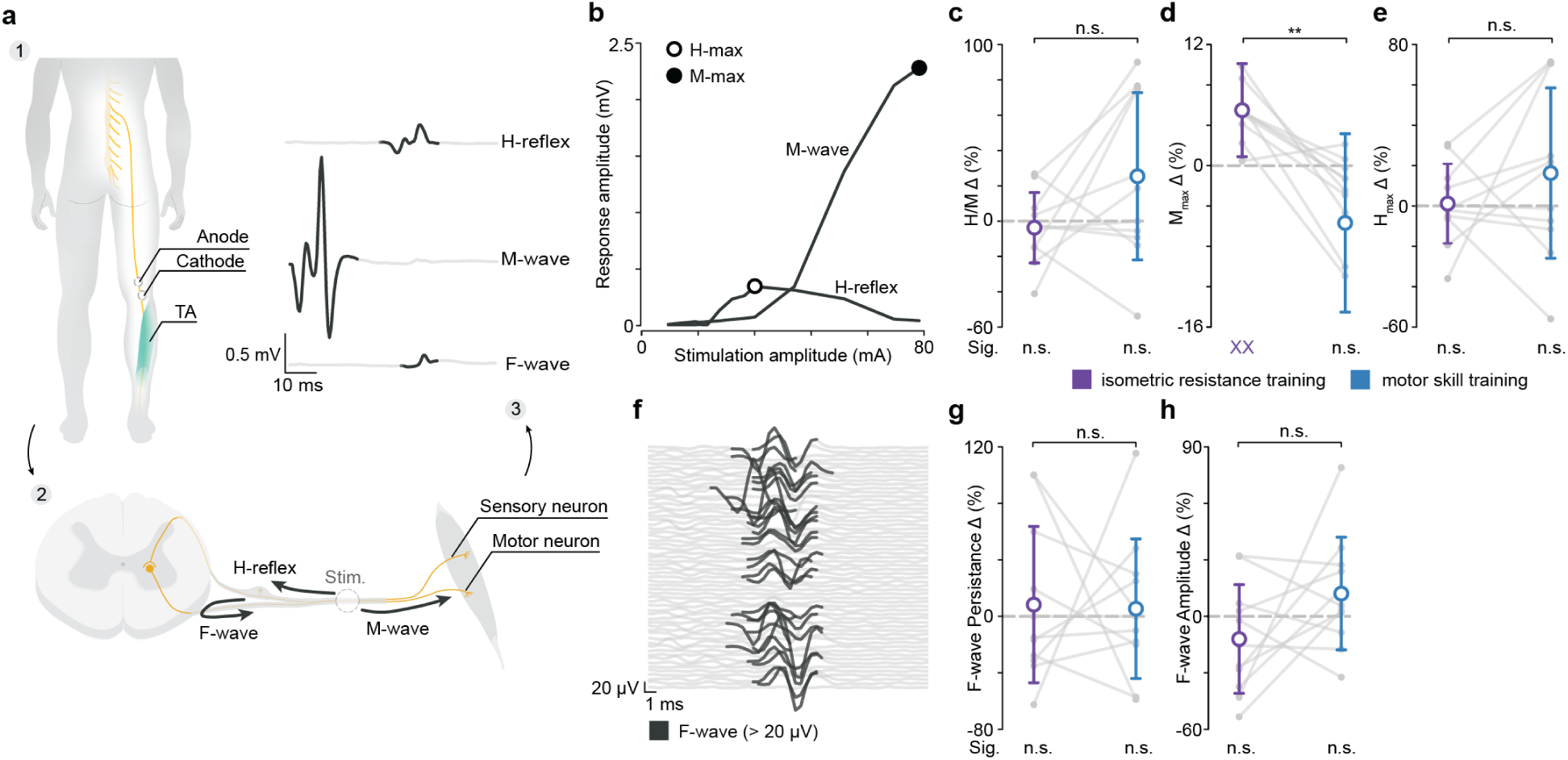
Effects of motor skill training and isometric resistance training on synaptic spinal excitability and spinal motor neuron excitability. (**a**) Peripheral nerve stimulation setup, response pathways, and representative responses. (1) Two 3.2 cm diameter surface electrodes were positioned on the popliteal fossa over the common peroneal nerve, with the anode placed 2 cm proximal to the cathode. (2) The M-wave arises from the direct activation of motor axons, leading to a short-latency response. The H-reflex follows stimulation of Ia afferent fibers, which synapse monosynaptically onto alpha motor neurons in the spinal cord, producing a long-latency, reflex-mediated response. The F-wave results from antidromic activation of alpha motor neurons, leading to depolarization of the soma and a recurrent orthodromic response via the same efferent motor pathway. (3) Representative H-reflex, M-wave, and F-wave responses are shown from participant MS018. The M-wave has been removed from the F-wave response for visualization purposes. (**b**) H-reflex and M-wave recruitment curves of participant MS018. (**c**) Percent change in H/M ratio. (**d**) Percent change in M_max_ amplitude. A significant increase in M_max_ was observed following isometric resistance training. (**e**) Percent change in H_max_. (**f**) Representative F-wave responses of participant MS014. (**g**) Percent change in F-wave persistence. (**h**) Percent change in F-wave amplitude. Asterisks above the group averages in (**c-e**) and (**g, h**) denote Bonferroni-corrected statistical significance from paired comparisons between training types: *p < 0.05, **p < 0.01. (Sig.) ‘X’ markers below the plot indicate Bonferroni-corrected significant training effects within each condition of percent change based on one-sample tests against µ = 0 (‘X’ p < 0.05, ‘XX’ p < 0.01, ‘n.s.’ p > 0.05).

We next sought to investigate training-induced changes in the maximum M-wave and H-reflex amplitudes (M_max_ and H_max_, respectively). M_max_ reflects the activation of muscle fibers via motor efferent axons and acts as a reference for normalizing reflex measures, yet it can either be potentiated or inhibited by exercise^52–54^. One-sample t-tests revealed significant increases in the M_max_ of the TA muscle after isometric resistance training (5.49% ± 4.61, t-statistic: −4.639, p-value: 0.002) but not after motor skill training (−5.67 % ± 8.84, t-statistic: 2.679, p-value: 0.064). Paired comparisons demonstrated that increases in M_max_ after isometric resistance training were significantly larger than after motor skill training (W = 1.00, p = 0.003) (**Figure 5d**). H_max_ is an indirect measure of the percentage of motor neurons that can be activated trans-synaptically^42,52,55^, although it can be influenced by factors such as presynaptic inhibition, afferent activity, and post-activation depression^42,52^. As with the H/M ratio, one-sample tests revealed that H_max_ did not significantly change after motor skill or resistance training (motor skill training: 16.31 % ± 41.85, W: 25.00, p-value: 1.0; resistance training: 1.20 % ± 19.51, t-statistic: 0.701, p-value: 1.0). Training-induced changes in H_max_ were also consistent across training paradigms (t-statistic: 1.083, p-value: 0.913) (**Figure 5e**). Together, our results suggest that while efferent fiber recruitment increases after isometric resistance training, this is not enough to impact the H/M ratio.

We used F-wave amplitude and persistence to compare changes in motoneuron excitability due to motor skill and resistance training^27,34,35^. F-waves are recurrent discharges of antidromically activated motoneurons, integrating both segmental and suprasegmental influences^27^. They appear at supramaximal stimulation amplitudes with variable latencies and amplitudes across repeated stimuli^27^ (**Figure 5f**). Individual one-sample tests revealed that F-wave persistence did not significantly increase after motor skill or resistance training (motor skill training: 5.39 % ± 49.40, t-statistic: −0.0354, p-value: 1.0; resistance training: −8.28 % ± 55.32 t-statistic: 0.973, p-value: 1.0). Training-induced changes in F-wave persistence were consistent across training paradigms (t-statistic: −0.1088, p-value: 1.0) (**Figure 5g**) F-wave amplitude similarly showed no significant changes after motor skill or resistance training (motor skill training: 12.11 % ± 29.90, t-statistic: −0.752, p-value: 1.0; resistance training: −12.06 % ± 28.87, t-statistic: 1.527, p-value: 0.473). Training-induced changes in F-wave amplitude were also consistent across training paradigms (t-statistic: 1.725, p-value: 0.346) (**Figure 5h**).

### Reticulospinal tract excitability remains consistent after motor skill and isometric resistance training

We used the StartReact response to evaluate reticulospinal tract excitability before and after training by comparing reaction times in response to visual, auditory, and startling stimuli (**Figure 6a,c**)^37^. The StartReact response refers to an accelerated release of a prepared motor action that is initiated by a startling stimulus^56,57^, and it has been used as a non-invasive method to assess the output gain of the reticulospinal tract, which operates through the auditory startle reflex (**Figure 6b**)^36,56^. Specifically, the reticulospinal tract gain (RST) is calculated by dividing the difference in reaction time between the startling stimulus and the visual stimulus by the difference in reaction time between the auditory stimulus and the visual stimulus (**Figure 6f**)^21,36^. We found a significant contribution of the reticulospinal tract for both motor skill (**Figure 6e**, pre: 1.69 ± 1.08 : W: 7.00, p: 0.009; post 1.21 ± 0.31 : W.00: 12.00, p: 0.034) and isometric resistance training (pre 1.29 ± 0.33 : t-statistic: 3.077, p: 0.042; post 1.28 ± 0.28 : t-statistic: 3.509, p: 0.020). However, one-sample tests showed no significant changes in RST gain after motor skill or isometric resistance training (**Figure 6f**, motor skill: −13.09% ± 30.00, W: 29.00, p-value: 0.459; isometric resistance: 4.52% ± 36.6, t-statistic: 0.058, p-value: 1.0). Furthermore, paired comparisons demonstrated consistency across training types (t-statistic: −1.313, p-value: 0.648). These results suggest that while there was a significant contribution from the reticulospinal tract for each task, the strength of this contribution was not impacted by training.

**Figure 6:**
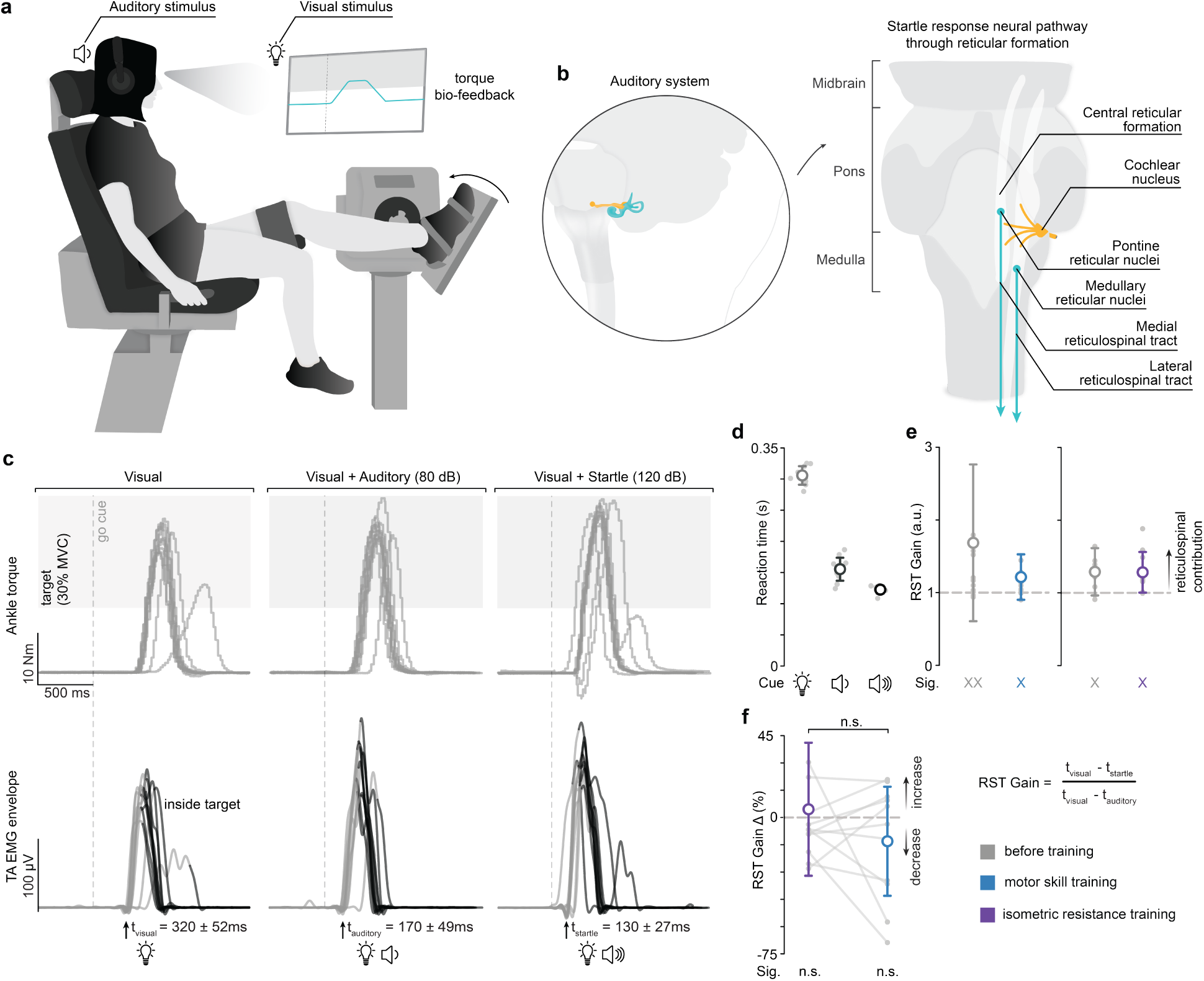
Reticulospinal tract excitability assessed via the StartReact paradigm remains unchanged following motor skill and resistance training. (**a**) StartReact paradigm to assess RST excitability. Participants sat in a Biodex chair and performed a rapid isometric dorsiflexion reaction task in response to visual and auditory cues. Real-time torque biofeedback was provided during testing to guide performance. (**b**). Schematic of the auditory startle pathway. The StartReact response assesses RST output gain non-invasively. Auditory signals from the cochlear nucleus activate giant neurons in the pontomedullary reticular formation, which then transmit signals via the RST to motoneurons for rapid motor responses. (**c**) Representative StartReact trial and RST gain calculation for representative participant ASR001. Torque (top) and tibialis anterior EMG profiles (bottom) show responses to each cue type: visual, visual + auditory (80 dB), and visual + startling stimulus (120 dB). The target contraction was set to 30% of MVC. (**d**) Individual reaction times from the trial shown in (**c**), plotted for each cue condition. Reaction times and variability decrease in response to the startling auditory cue, reflecting a startle response. (**e**) Reticulospinal tract gain. Group-level RST gain values before and after motor skill and resistance training. ‘X’ marker below plots indicates Bonferroni-corrected significant contribution of the RST at each timepoint based on one-sample tests against µ = 1 (‘X’ p < 0.05, ‘XX’ p < 0.01, ‘n.s.’ p > 0.05). (**f**) Training-induced percent changes in RST gain. Neither training protocol measurably altered the influence of the reticulospinal tract on the rapid initiation of a prepared movement in response to a startling stimulus.

## Discussion

### Motor and resistance training increased corticospinal tract excitability, with minimal spinal or reticulospinal circuit changes

Our findings demonstrate distinct short-term adaptations in corticospinal tract excitability following single sessions of cue-paced motor skill and cue-paced isometric resistance training. Contrary to our initial hypothesis, both training paradigms enhanced corticospinal tract excitability, albeit to differing degrees, while neither modality induced measurable changes in reticulospinal or spinal motoneuron excitability. MEP amplitude may increase to different extents in lower limb muscles depending on training type, measurement modality, and excitability in both the cortical and spinal motor neuron pool^50^.

The robust increase in corticospinal tract excitability following motor skill training (**Figure 4**) aligns well with existing evidence indicating that skill-dependent training is particularly effective in inducing corticospinal plasticity^13,58,59^. This effect was notably more pronounced at rest compared to muscle pre-activation. Our results are in agreement with previous studies showing that MEP amplitudes are generally higher during pre-activation of the targeted muscle^60,61^. These findings contradict a previous meta-analysis by Siddique et al., showing that resistance training increased MEP amplitudes during voluntary contractions but not at rest^30^. We found that pre-activation reduces within-participant variability, making it a useful tool for accurately and precisely evaluating corticospinal tract excitability at a single time point. Yet, the relatively small change in MEP amplitudes during the pre-activation condition compared to rest suggests that this reduction in variability may obscure differences in corticospinal tract excitability across evaluations. Therefore, we recommend carefully considering whether TMS evaluations should be performed at rest or during pre-activation, as our findings highlight the importance of selecting the appropriate muscle activation state based on whether the goal is to maximize signal detection or assess task-specific adaptations.

Our assessment of spinal excitability through the H-reflex, M-wave, and F-wave metrics showed minimal training-induced changes (**Figure 5**). This aligns with previous findings suggesting that significant spinal plasticity typically requires sustained, high-intensity, or repetitive training paradigms^35,52,53^. Notably, we observed an increase in the M_max_ following isometric resistance training, suggesting that this training paradigm influenced peripheral neuromuscular properties, such as increased motoneuron recruitment, in addition to changes in central nervous system excitability.

These observations echo previous findings reporting increases in corticospinal tract excitability following 4 weeks of visuomotor skill learning but not heavy strength training^62^. Similarly to our findings, no substantial short-term changes in spinal excitability were observed, suggesting central nervous system plasticity primarily occurred at higher cortical rather than spinal levels. However, Jensen et al. observed a decrease in corticospinal tract excitability after prolonged strength training, while our study found an increase after a single session. One potential reason for this discrepancy is that Jensen et al. targeted upper limb muscles (biceps brachii), whereas our study focused on lower limb muscles (tibialis anterior). Therefore, differential involvement of neural tracts (corticospinal vs. reticulospinal) due to anatomical and functional distinctions between upper and lower limb movements could underline differences in neural excitability outcomes.

### Lack of increase in reticulospinal tract excitability following distal motor training may reflect insufficient engagement or limitations in StartReact paradigm sensitivity

The StartReact response has been used as a non-invasive method to assess the output gain of the reticulospinal tract, which operates through the auditory startle reflex^36,56^. The auditory pathway, beginning at the cochlear nucleus, directly activates giant neurons in the pontomedullary reticular formation^57^. Following this activation, the signal is transmitted to motoneurons in the brainstem and spinal cord via the reticulospinal tract, facilitating rapid motor responses^57^. Although the StartReact paradigm effectively measured a significant contribution from the reticulospinal tract during the assessed tasks (**Fig. 6e**), neither motor skill nor the isometric resistance training induced changes in reticulospinal tract excitability (**Fig. 6f**). Given previous reports highlighting reticulospinal involvement in gross motor tasks, coordinated finger movements, and strength generation^21,56,63^, our findings have several possible explanations. First, as the reticulospinal tract is thought to contribute more to balance and strength production^20,22^, the distal joint nature of our training tasks may inherently limit reticulospinal engagement. Second, while the reticulospinal gain is consistent across upper limb muscles in unimpaired individuals, it has been shown to be heightened in people with SCI in extensor muscles compared to flexor muscles^64^ or in the case of spasticity^63^. Therefore, it remains possible that training-induced changes in reticulospinal tract contributions could be different for people with damage to the central nervous system, such as in SCI. Third, the lack of measurable changes by any training intervention may suggest that the StartReact response is not sufficiently sensitive to detect changes in the excitability of the reticulospinal tract or that a single session is not sufficient to induce these changes. Fourth, changes in excitability after single training sessions may be less likely in subcortical structures compared to the cortex.

### Reticulospinal sparing and corticospinal plasticity may have important clinical implications for targeted neurorehabilitation

Biological repair strategies may traditionally focus on regenerating corticospinal projections^65–67^ because of their high contribution to dexterous tasks in humans^68,69^. However, reticulospinal circuits contribute to recovery after corticospinal tract lesions or SCI^21,70^, and their wide distribution in the white matter^71,72^ and direct and indirect projections to spinal motor neurons^73^ make reticulospinal circuits prone to be partially spared after SCI in humans^74^. Moreover, reticulospinal tract neurons can display equal or greater responses to regeneration strategies after injury compared to corticospinal tract axons^75–77^, leading to more potential for recovery after SCI^78^, despite reticulospinal sparing being associated with increased spasticity^63^. Our findings suggest that single session motor skill and isometric resistance training protocols may be insufficient to engage the reticulospinal tract in the short term, highlighting a possible limitation of these approaches. Studies in SCI populations or across extended rehabilitation timelines could provide valuable insights into how to more effectively recruit reticulospinal pathways through training.

Additionally, skill-based training should be prioritized in rehabilitation programs targeting corticospinal plasticity^58^. Individuals with limited mobility can still perform skill-based training, potentially benefiting from its neuroplastic effects. Our findings enhance the understanding of motor control and neural plasticity, supporting the emerging perspective that rehabilitation strategies should target specific neural pathways according to functional goals^28,79,80^. Examining variations in training intensity, duration, and task complexity will help identify optimal interventions for promoting neural plasticity. Longitudinal studies are crucial for evaluating long-term adaptations. Furthermore, integrating neurophysiological measures with functional assessments—such as gait analysis—could significantly enhance translational potential and clinical applicability. In conclusion, while our study provides valuable insights into the effects of motor skill and resistance training on corticospinal, reticulospinal, and spinal excitability, future work is needed to refine training protocols and assessment methods to maximize rehabilitation outcomes.

## LIMITATIONS OF STUDY

Our study advances previous research by concurrently examining multiple neural substrates involved in motor control—namely corticospinal, reticulospinal, and spinal circuits—in response to distinct training paradigms. This comprehensive approach provides broader insight into neural adaptations and interactions across different levels of the nervous system. However, the relatively small sample size (N = 23) and homogeneous population (26-year-old average unimpaired participants) restrict the generalizability to clinical populations.

Although motor-evoked potentials are a valuable non-invasive measure of corticospinal tract excitability^81^, they do not reflect intracortical excitability or inhibition, which may be important contributors to training-induced changes. Future studies employing paired-pulse TMS techniques to evaluate short-interval intracortical inhibition (SICI) and facilitation (ICF) could further reveal specific intracortical processes underlying training-induced changes in excitability^82,83^.

While short-term changes in corticospinal and intracortical excitability remain present 75 min after the intervention^31^, it is unclear how long they remain stable. Therefore, increasing the number of evaluations and, thus, the experiment time could play a role in the potential observed effects. It is important to note that although training time was limited to 30 min, experimental sessions lasted approximately 2.5 to 3 hours due to EMG preparation time (∼30 min) and neurophysiological evaluations (∼1 hour before training and ∼1 hour after training). Therefore, although evaluating neural excitability on additional pathways, muscles, time points, and under diverse conditions would certainly improve the clarity of changes in excitability^31,35,37^, it is crucial to consider participant comfort and avoid excessive session durations, as the 3-hour sessions could already approach the practical limits of participant endurance, attention, and retention.

## RESOURCE AVAILABILITY

### Lead Contact

Further information and requests for resources should be directed to and will be fulfilled upon reasonable request by the lead contact, Dr. Ismael Seáñez.

### Data and code availability

- Data from this study will be made available upon reasonable request to the lead contact.
- All software used to produce the figures in this manuscript will be made available upon reasonable request to the lead contact.
- Any additional information required to reanalyze the data is available from the lead contact upon request.

## ACKNOWLEDGEMENTS

This work was supported in part by the National Institutes of Health NINDS Award Number K01NS127936. The NICHD Award Number K12HD073945, and internal funding from the Department of Biomedical Engineering, the Department of Neurosurgery, and the McDonnell Center for Systems Neuroscience at Washington University in St. Louis provided additional support. R.H., C.A., R.K., Z.S., H.N., N.P., and I.S., received partial support from these sources.

## AUTHOR CONTRIBUTIONS

R.H., N.P., and Z.S. collected the data.

R.H. and N.P. analyzed the data.

R.K., Z.S., and H.N., developed software to run experiments.

R.H., C.A., R.K., and I.S., contributed to study design and data interpretation.

R.H. and I.S. wrote the manuscript.

I.S. conceptualization and supervision.

## DECLARATION OF GENERATIVE AI AND AI-ASSISTED TECHNOLOGIES IN THE WRITING PROCESS

During the preparation of this work, the author(s) used ChatGPT AI and Grammarly AI for grammatical editing purposes. After using this tool or service, the author(s) reviewed and edited the content as needed and take(s) full responsibility for the content of the publication.

## STAR★METHODS

### EXPERIMENTAL MODEL AND STUDY PARTICIPANT DETAILS

#### Participants

Thirty-two unimpaired participants provided informed consent to participate in this study, which was reviewed and approved by Washington University in St. Louis’ Institutional Review Board, and all 32 began participation in the experiments. Nine individuals were excluded from analysis for the following reasons: protocol change (N=3), participant personal scheduling issue (N=1), TMS motor threshold too high (N=1), could not tolerate TMS (N=1), and identified as outliers (N=3; one due to abnormal MEP exceeding 100% M_max_, two due to EMG recording errors affecting reaction time measurement). Participant demographics, group designation, and excluded participants are listed in **Supplementary Table S1.**

The study was divided into two distinct groups. Group A, consisting of 11 participants (6 females and 5 males, with an average age of 26.18 ± 4.99 years), performed corticospinal and spinal excitability evaluations before and after training. Group B, comprising 12 participants (7 females and 5 males, with an average age of 26.92 ± 7.05 years), performed reticulospinal tract excitability evaluations before and after training. Both groups underwent the same cross-over design, which included a 30-minute motor skill training session and a 30-minute isometric resistance training session on separate days (**Figure 1**). The order of these sessions was randomized for each participant.

### METHOD DETAILS

#### Motor skill training

During motor skill training, participants sat in a Biodex (System 4 Pro™, Biodex Medical Systems, USA) isokinetic dynamometer, with both legs hanging freely. They used a body-machine interface (BoMI) to control a computer cursor (**Figure 2a**). The BoMI comprised of four non-invasive, wireless inertial measurement units (IMUs) (3-Space™ Sensors, Yost Labs, USA) secured to adjustable straps on the participant’s right foot to record foot and ankle movements.

During the calibration phase, participants were instructed to perform "free movements" within a comfortable range for 55 seconds. A PCA was performed on the Euler angles obtained from the IMUs to identify the principal components that accounted for the greatest amounts of variance^43^. The top two principal components were used to control the horizontal and vertical movements of a computer cursor so that the 8-dimensional body motion vector (roll and pitch from 4 sensors) was projected into a 2-dimensional cursor control vector (cursor x and y). Participants could then use their ankle movements to control the computer cursor.

After calibration, participants familiarized themselves with the BoMI before completing five center-out-reaching tasks over the 30-minute session^44–47^. Participants wore headphones (U UFO Over-Ear Headphones, Bluedio, China), which provided a 1 Hz auditory cue to guide them to time their target reaches to conclude at four seconds, encouraging slow and steady movements. If the tasks were completed before the 30-minute mark, participants engaged in a maze game of increasing difficulty to maintain consistent training duration across training paradigms.

#### Isometric resistance training

In the isometric resistance training session (**Figure 3a**), participants sat in the Biodex chair with the left leg resting on a footrest and the right leg positioned with the foot attached to the dynamometer. The hip was positioned at 120°, the knee at 160°, and the ankle at 110° (measured from neutral), with a permissible variance of ±10° to ensure participant comfort and prevent muscle stretch^84^. Before the training began, the maximum torque for plantar flexion and dorsiflexion was obtained by asking participants to perform three consecutive maximal effort ankle joint pushes (for plantar flexion) or pulls (for dorsiflexion). During the training, participants executed slow-ramped movements to reach 30% of their maximum torque in both directions. The training comprised of three blocks of 12 contraction trials, each trial including both dorsiflexion and plantar flexion isometric contractions, for a total training time of 30 minutes. Participants took a 30-second rest between trials and a 2-minute break between blocks. To help maintain consistent velocity during the ramped movements, a 1 Hz auditory cue was provided, similar to the one used in motor skill training. Participants used the cue to guide a sequence of 3-second dorsiflexion and plantarflexion ramps to 30% of their maximum voluntary contraction (MVC), each followed by 3-second holds and returns to rest. The auditory cue provided clear timing for when to initiate, sustain, and release each contraction.

#### Electromyography data acquisition

Wireless surface electrodes (Trigno® Avanti, Delsys Inc., USA) were used to record electromyography (EMG) data with sensors placed bilaterally according to SENIAM guidelines covering the rectus femoris, vastus lateralis, tibialis anterior (TA), medial gastrocnemius (MG), and soleus. Prior to electrode placement, the skin over the muscle belly was shaved, if necessary, and cleaned using abrasive gel (NuPrep®, Weaver and Co., USA) applied with a Q-tip®, followed by wiping with alcohol pads. An additional EMG sensor with analog input (DC-X06 Analog Input, Delsys Inc., USA) was used to capture stimulation onset for offline alignment between stimulation pulses and EMG data. EMG data was amplified using a data acquisition system (Trigno® Avanti Research+, Delsys Inc., USA; gain: 300; bandwidth 20–450 Hz), and sampled at either 2148 Hz or 2000 Hz, with the difference in sampling rate due to a lab software update that occurred mid-study. All EMG data was displayed in real-time using custom-built software written by our group in Python v3.11^85^.

#### Transcranial magnetic stimulation

TMS targeting the left motor cortex was used to evoke motor-evoked potentials (MEPs) in the right tibialis anterior to assess the excitability of the corticospinal tract^13,49^ (**Figure 4a**). A Magstim 200^2^ (MOP01-EN, Magstim, UK) stimulator with a 110 mm double-cone coil (MOP21-EN^86^, peak magnetic field of ≥ 1.2T) and a monophasic (100 ms) waveform was employed for TMS (**Figure 4b**). Participants sat in the Biodex isokinetic dynamometer with their hips positioned at 120°, knee at 160°, and ankle at 110°, with the right foot strapped securely to the dynamometer to ensure consistent positioning during stimulation. The Cz point, representing the scalp vertex, was identified by finding the intersection between the line from the nasion to the inion and the line between the left and right pre-auricular points. The intersection point, Cz, was marked using a skin-safe marker (Tondaus®, Model T3023, Surgical skin marker, China).

The TMS coil and participant’s head 3D position and orientation were tracked via infrared cameras (Miqus Hybrid, Qualysis, Sweden) and reflective markers (Super-spherical Markers, Qualysis, Sweden) using a custom-built NeuroNavigation software designed in MATLAB (2024a, Mathworks, USA) (**Figure 4b**). At the start of each session, a large grid search was performed to identify the hotspot for the right TA muscle, which is approximately 1.6 cm lateral and 0.8 cm posterior to Cz^87^. The TMS coil was positioned at a 45° angle to induce postero-anterior (PA) current flow^88^. Stimulation intensity was gradually increased from 30% maximum stimulator output at 5% increments until a short-latency response (20-30 ms) was observed in the contralateral TA with a peak-to-peak amplitude of at least 50 µV. The stimulator intensity was recorded as the general motor threshold, and the TMS coil 3D position and orientation with respect to the participant’s head were saved as the initial hotspot location for the TA. A hotspot search was then conducted using 120% of the general motor threshold to identify the optimal location for TA targeting based on the largest evoked MEPs. MEPs were evoked at 9 locations spaced 1 cm apart following a 3×3 grid pattern around the initial hotspot, while the peak-to-peak amplitudes of the MEPs were quantified and displayed in real-time using the custom-built software. A higher-precision search was conducted around the location with the highest MEP amplitude by moving the coil 0.3 cm in the posterior, anterior, medial, and lateral directions. The 3D position and head orientation with the highest MEP amplitude were saved as the optimum TA hotspot, and the resting motor threshold was determined at this location.

To find the TA resting motor threshold at the identified hotspot, the stimulator intensity was reduced in 1-2% increments until the lowest intensity that could evoke an MEP in at least 3 out of 5 stimuli with a peak-to-peak amplitude ≥ 50 µV^13,58,89,90^. Recruitment curves for MEP responses (**Figure 4c**) were collected at intensities of 100%, 150%, 180%, and 200% of the resting motor threshold presented in a randomized order with ten repetitions for each intensity. These recruitment curves were obtained before and after training, with the higher-precision search conducted post-training around the hotspot identified before training (**Figure 4d**). TMS evaluations were conducted both at rest and during 15% dorsiflexion contraction using a custom-built real-time visualization software displaying the moving average of TA EMG feedback (Python v3.11). In the pre-activation condition, the experimenter manually delivered the TMS trigger as soon as the participant’s TA EMG exceeded the 15% MVC threshold target. Using EMG allows for a more consistent evaluation of neuromuscular drive, even if participant force production declines due to fatigue after training^91^. Participants were able to rest between pulses, and the order of active vs. passive evaluations was randomized for each participant. As some participants could not tolerate all TMS intensities, the number of samples varied across TMS intensities.

#### Peripheral nerve stimulation

Peripheral nerve stimulation targeting the posterior tibial and common peroneal nerves was used to assess spinal synaptic transmission efficacy of Ia sensory fibers to the homonymous motor fibers^51,52^. Supramaximal electrical stimulation was used to assess alpha-motoneuron excitability through the antidromic activation of motor neurons, serving as a complementary method to evaluate the lower neuronal circuits^27,41^ (**Figure 5a**). Stimulation was delivered using a biphasic constant-current stimulator (Digitimer DSR8, Digitimer Ltd., U.K.) with round surface electrodes (PALS Neurostimulation Electrodes, Axelgaard Manufacturing Co., Ltd., USA) of a 3.2 cm diameter and a National Instruments USB-6001 (NI USB 6001, National Instruments, USA) to trigger 1 ms (per phase) biphasic stimulation pulses. The stimulating cathode was placed on the popliteal fossa of the right leg, with the return anode positioned 2 cm proximal to the cathode (**Figure 5a**). The electrode configuration allowed for the elicitation of H-reflex, M-waves, and F-waves from the MG and TA simultaneously^92,93^. A recruitment curve was created using a non-linear sweep of sixteen stimulation amplitudes, ranging from 50% of the lowest H-reflex threshold to 110% of the highest stimulation amplitude that produced the maximal M-wave response (M_max_)^94^ (**Figure 5b**). Three repetitions were performed at each stimulation amplitude with a 7-second interval between pulses. F-waves were elicited by delivering 60 pulses at 125% of M_max_ amplitude, with a 1-second interval between pulses (**Figure 5f**).

#### StartReact evaluation

StartReact refers to an accelerated release of a prepared motor action that is initiated by a startling stimulus^56,57^. This phenomenon occurs when a startling stimulus is introduced during the preparation phase of a voluntary movement, resulting in the execution of the planned action occurring significantly faster than non-startling auditory or visual cues^57^. A custom-built software (Python v3.11) was used to collect synchronized torque and EMG data stream and display them as biofeedback in real-time (43-inch Class FULL HD LCD Display, Panasonic, Japan). During the StartReact task, participants sat in the Biodex (System 4 Pro™, Biodex Medical Systems, USA) as described above with their right foot secured to the ankle attachment plate (**Figure 6a**). Headphones rated at 120 dB SPL (U UFO Over-Ear Headphones, Bluedio, China) were connected to an amplifier (AV Surround Receiver AVR-S510BT, Denon, Japan) and used to deliver auditory cues. Upon receiving a ‘go’ cue, participants were instructed to perform an isometric contraction by dorsiflexing their right ankle as quickly as possible. The cue types included a visual cue, a visual + auditory cue (80 dB, 500 Hz, 50 ms), and a visual + startle cue (120 dB, 500 Hz, 50 ms)^37^. The contraction target was set to 30% of their MVC (**Figure 6a**). Ten trials of each cue type were performed in randomized order for a total of 30 trials per block, with 1-3 second rest periods (randomized time) between cues.

### QUANTIFICATION AND STATISTICAL ANALYSIS

#### Motor learning

Motor learning was assessed by comparing reaching performance in center-out tasks using several performance metrics: path length, movement time, jerk, and endpoint error^45^. Path length refers to the total distance traveled by the cursor during the center-out reaching tasks. Movement time is the duration from the initiation of movement at the center target until the final target is reached. Dimensional jerk, which quantifies movement smoothness, is calculated from the third derivative of position^95^. Endpoint error is defined as the Euclidean distance between the final cursor position and the target location after four seconds. Performance metrics were averaged across 24 movements per block after outlier detection and removal using the interquartile range (IQR) method, defined as values exceeding 1.5 times the IQR above the 75th percentile or below the 25th percentile^16,53^. Motor learning was quantified by comparing improvements in movement performance from the first to the last block after the 30-min training session.

#### Torque and muscle capacity before and after training

Participants were instructed to perform three repetitions of maximal voluntary contraction to assess muscle capacity before and after training. The EMG signal was first full-wave rectified by removing the mean and taking the absolute value. The root mean square (RMS) of the rectified signal was then calculated using a sliding window of 100 ms. The onset and offset of muscle activation were established by detecting instances when the RMS amplitude surpassed a manually set threshold, typically around 0.02 mV. Manual threshold selection was incorporated into the data analysis to address baseline signal variability, ensuring the robustness of burst identification for all individuals. The area under the curve (AUC) for each burst was computed using the trapezoidal rule, applied to the RMS signal between the identified onset to 2.25 seconds later (**Figure 3c**). The muscle capacity was quantified as the average AUC across the three MVC bursts recorded before and after the training.

#### Corticospinal tract excitability

Corticospinal tract excitability was evaluated by examining changes in MEP amplitude from baseline to post-training^53,58,86^. MEP peak-to-peak amplitudes were normalized to the maximal M-wave amplitude (M_max_) recorded from peripheral nerve stimulation at the corresponding time point (pre- or post-training), with responses expressed as a percentage of M_max_^13,53^. Following normalization, values that exceeded 100% were excluded, as M_max_ represents the excitation of the entire motoneuron pool^52^, making MEPs (% of M_max_) larger than 100% unreasonable and indicative of a potential recording error. Outliers were identified and removed using the interquartile range (IQR)^16,53^. The remaining normalized MEP percentages were averaged across trials for each TMS intensity. Changes in normalized MEP amplitudes following training were computed for each TMS intensity by comparing pre- and post-training averages using the formula:

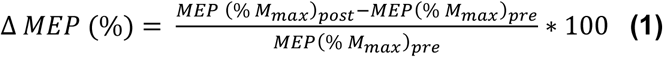

#### Synaptic and spinal motor neuron excitability

The H/M ratio was used to assess synaptic spinal excitability before and after training^32,33^. This ratio serves as an estimate of the excitability of the spinal reflex pathway by evaluating the activation of the motor neuron pool via the Ia afferents, in relation to the maximum capacity of the entire motor neuron pool^42,52,55^. The peak-to-peak amplitudes of the M-wave and H-reflex responses were averaged across the three repetitions at each stimulation level. This process created a recruitment curve for both the M-wave and H-reflex (**Figure 5b**). H_max_ and M_max_ values were identified from this curve and were used to calculate the H/M ratio. To assess training effects, the percent change in the H/M ratio from pre- to post-training was computed for each participant using an analogous form of Equation 1.

F-wave amplitude and persistence were used to assess spinal motor neuron excitability^27,34,35^. EMGs were filtered using a second order Bessel high-pass filter (200 Hz) to improve the clarity of F-wave evaluations^81^. An F-wave response was deemed valid if it occurred at least 30 ms after the stimulus and had a peak-to-peak amplitude greater than or equal to 20 µV^81^. F-wave persistence was calculated as the percentage of stimuli out of 60 pulses that elicited a valid response. To assess training effects, the percent change in average amplitude and persistence from pre- to post-training was computed for each participant using an analogous form of Equation 1^35^.

#### Reticulospinal tract excitability

Reticulospinal tract (RST) excitability was evaluated by comparing the StartReact response before and after training^37^. The StartReact response, or RST gain, is calculated as follows (**Figure 6c**):

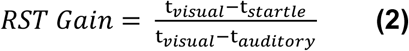

The ratio is the difference in reaction time between the startling (t_startle_) and visual cues (t_visual_) divided by the difference in reaction time between the auditory (t_auditory_) and visual cues^21,36^. Reaction time was defined as the time between cue onset and movement onset. Movement onset was identified from EMG traces (**Figure 6c**) offline using a Python library for change point detection (ruptures v1.1.9), specifically employing a binary segmentation search method (model = “l2”, Binseg method, predict 8 changepoints)^96^. Reaction times less than 75 ms were excluded as false starts, and reaction times greater than 700 ms were discarded as indicating a loss of participant focus^37^. Additionally, reaction times identified as outliers based on the IQR were excluded from analysis. To assess training effects, the percent change in RST Gain from pre- to post-training was computed for each participant.

#### Statistics

Statistical analyses were performed using Python (v3.11) using the SciPy and StatsModels libraries. To assess the normality of all data sets, the Shapiro-Wilk test was applied. Given the non-normal distribution of several datasets, both parametric and non-parametric statistical tests were used, with significance set at p < 0.05. Group data are reported as mean ± standard deviation (SD), unless stated otherwise.

To test for motor learning in the BoMI, paired t-tests (for normally distributed data) and Wilcoxon signed-rank tests (for non-normally distributed data) between the first and last blocks of training were conducted under the null hypothesis that there would be no significant reduction in jerk, movement time, or path length following motor skill training (i.e. no motor learning occurred). To assess neuromuscular fatigue, paired t-tests (for normally distributed data) and Wilcoxon signed-rank tests (for non-normally distributed data) were used to compare pre- and post-training MVC EMG amplitudes. The null hypothesis was that there would be no difference in EMG amplitude before and after training (i.e. neither training would induce a reduction in EMG amplitude). These tests also assessed whether the degree of fatigue, indicated by changes in EMG amplitude, differed between the two training modalities, with the null hypothesis stating no difference in training effects.

A Generalized Linear Mixed Model (GLMM) was used to analyze changes in corticospinal excitability, considering the unequal number of data points at each TMS amplitude. This model investigated the effects of training conditions (motor skill training vs. isometric resistance training), pre-activation states (resting vs. active), and TMS intensity on the percent change in motor-evoked potential (MEP) amplitude, treating percent change in MEP amplitude as the dependent variable. Training conditions and pre-activation states were considered fixed categorical variables, while TMS intensity was regarded as a fixed continuous variable. The participant was included as a random effect. If the GLMM indicated a significant increase in excitability due to a protocol, paired t-tests (for normally distributed data) and Wilcoxon signed-rank tests (for non-normally distributed data) with a Bonferroni correction for multiple comparisons were used to compare the effect between training conditions and one-sample tests were used to evaluate individual training/activation conditions to reject the null hypothesis that the average change in MEP was equal to 0.

All statistical analyses for reticulospinal, synaptic spinal, and spinal motor neuron excitability followed a standardized workflow. Data were first assessed for normality using the Shapiro–Wilk test. Depending on normality, either paired t-tests or Wilcoxon signed-rank tests (with Bonferroni correction for multiple comparisons) were used to compare measurements between the two training conditions. These tests evaluated whether one training modality elicited significantly greater or lesser changes in excitability relative to the other, with the null hypothesis being that there was no difference between training effects. Additionally, one-sample t-tests or Wilcoxon signed-rank tests were applied to assess within-condition percent changes from baseline. For these, the null hypothesis was that the mean percent change was equal to zero, indicating no change in excitability following training. Corrected effect sizes were calculated to complement inferential statistics with Hedge’s g to account for the small sample size bias^97^.

## SUPPLEMENTAL INFORMATION

Document S1. Table S1 and S2

Table S1: Participant demographics

Table S2: GLMM results of corticospinal tract excitability

