## Supplemental Tables for "Transient effects in corticospinal and reticulospinal tract excitability induced by motor skill and isometric resistance training"

| ID | Age (y) | Sex | Height (m) | Weight (kg) | Race | Hispanic/ Latino | Experiment | |
|---|---|---|---|---|---|---|---|---|
| | | | | | | | Cortical & Spinal | Reticulospinal |
| MS001 | 26 | Male | 1.63 | 58.97 | Asian | No | ✓ | |
| MS002 | 23 | Female | 1.65 | 54.43 | White | No | ✓ | |
| MS003 | 28 | Male | 1.88 | 83.91 | Asian, White | No | Protocol change | |
| MS004 | 23 | Female | 1.65 | 62.14 | Asian, White | No | ✓ | |
| MS005 | 25 | Female | 1.60 | 54.00 | Asian | N/A | Protocol change | |
| MS006 | 24 | Male | 1.78 | 63.5 | White | No | ✓ | |
| MS007 | 23 | Male | 1.78 | 106.59 | White | No | Protocol change | |
| MS008 | 23 | Female | 1.60 | 49.90 | Asian | No | ✓ | |
| MS009 | 54 | Female | 1.55 | 74.84 | White | No | Did not return for follow-up | |
| MS010 | 22 | Female | 1.80 | 68.04 | White | No | ✓ | |
| MS011 | 27 | Male | 1.83 | 88.45 | White | No | Withdrawn – TMS thresholds too high | |
| MS012 | 39 | Female | 1.70 | 65.77 | White | No | ✓ | |
| MS013 | 23 | Female | 1.65 | 61.24 | White | Yes | Identified as outlier | |
| MS014 | 28 | Male | 1.73 | 81.65 | White | No | ✓ | |
| MS015 | 28 | Female | 1.75 | 70.31 | American Indian or Alaskan Native | Yes | Withdrew – could not tolerate TMS | |
| MS016 | 31 | Female | 1.63 | 62.60 | Asian | No | ✓ | |
| MS017 | 28 | Male | 1.73 | 53.00 | Asian | No | ✓ | |
| MS018 | 21 | Male | 1.96 | 102.06 | Asian, White | Yes | ✓ | |
| ASR001 | 29 | Male | 1.70 | 79.38 | White | No | | ✓ |
| ASR002 | 26 | Male | 1.63 | 61.24 | Asian | No | | ✓ |
| ASR003 | 23 | Female | 1.65 | 52.16 | White | No | | ✓ |
| ASR004 | 27 | Female | 1.73 | 61.24 | White | No | | ✓ |
| ASR005 | 20 | Male | 1.80 | 69.85 | Asian, White | No | | Identified as outlier |

| | | | | | | | |
|---|---|---|---|---|---|---|---|
| ASR006 | 21 | Male | 1.57 | 65.00 | Asian | No | ✓ |
| ASR007 | 22 | Female | 1.80 | 68.04 | White | No | ✓ |
| ASR008 | 28 | Male | 1.83 | 80.74 | White | No | Identified as outlier |
| ASR009 | 28 | Female | 1.75 | 68.04 | American Indian or Alaska Native | Yes | ✓ |
| ASR010 | 23 | Female | 1.65 | 61.24 | White | Yes | ✓ |
| ASR011 | 25 | Female | 1.73 | 95.25 | White | No | ✓ |
| ASR012 | 26 | Male | 1.96 | 90.72 | Black or African American | No | ✓ |
| ASR013 | 49 | Female | 1.65 | 61.24 | White | No | ✓ |
| ASR014 | 24 | Male | 1.91 | 83.91 | White | No | ✓ |

**Table S1.** Participant demographics information for the neurophysiology experiments. Participants who withdrew or were excluded from the analysis do not have a checkmark, and the reason for exclusion is specified instead.

| | | | | | | |
|---|---|---|---|---|---|---|
| No. Observations: | 178 | Method: | REML | | | |
| No. Groups: | 12 | Scale: | 1327.5949 | | | |
| Min. group size: | 12 | Log-likelihood: | -892.7539 | Converged: | Yes | |
| Max. group size: | 16 | Mean group size. | 14.8 | | | |
| | | | | | Confidence Interval | |
| Effects | Coef. | Std. Err. | z | P>\|z\| | [0.025 | 0.975] |
| motor skill training | 37.213 | 13.2 | 2.819 | 0.005 | 11.342 | 63.084 |
| isometric resistance training | 28.554 | 13.253 | 2.154 | 0.031 | 2.578 | 54.530 |
| Pre-activation | -17.656 | 5.479 | -3.222 | 0.001 | -28.395 | -6.917 |
| % RMT | -0.060 | 0.074 | -0.811 | 0.417 | -0.204 | 0.085 |
| Group Var. | 271.673 | 4.439 | | | | |

**Table S2.** GLMM results (3 factors: TMS intensity (% RMT) [100%, 150%, 180%, 200]; training [motor skill training vs. isometric resistance training]; evaluation condition [rest vs. pre-activation (15% dorsiflexion)]). Results related to **Fig. 4e,f**.
